# Disentangling the neurobiological bases of temporal impulsivity in Huntington’s disease

**DOI:** 10.1101/2023.02.07.527472

**Authors:** Helena Pardina-Torner, Audrey E. De Paepe, Clara Garcia-Gorro, Nadia Rodriguez-Dechicha, Matilde Calopa, Jesus Ruiz-Idiago, Celia Mareca, Ruth de Diego-Balaguer, Estela Camara

**Author notes:** <u>Corresponding author: Estela Camara</u>, Cognition and Brain Plasticity Unit, IDIBELL (Institut d’Investigació Biomèdica de Bellvitge), Feixa Larga S/N, 08907 L’Hospitalet de Llobregat (Barcelona), Spain, www.brainvitge.org.

## Abstract

**Background:** Despite its impact on daily life, impulsivity in Huntington’s disease (HD) is understudied as a neuropsychiatric symptom. Our aim is to characterize temporal impulsivity in HD, evaluated through a Delay Discounting (DD) task, and to disentangle the underlying white matter correlates in HD.

**Methods:** Forty-seven HD individuals and thirty-six healthy controls conducted a DD task and complementary Sensitivity to Punishment and Sensitivity to Reward (SR) Questionnaire. Diffusion-tensor imaging was employed to characterize the structural connectivity of two limbic tracts: the uncinate fasciculus (UF) and the accumbofrontal tract (NAcc-OFC). Multiple linear regression analyses were applied to analyze the relationship between impulsive behavior and white-matter microstructural integrity.

**Results:** Altered structural connectivity in both the NAcc-OFC and UF in HD individuals was observed. Moreover, the variability in structural connectivity of these tracts was associated with the individual differences in temporal impulsivity. Specifically, increased structural connectivity in the right NAcc-OFC predicted increased temporal impulsivity, while reduced connectivity in the left UF was associated with higher temporal impulsivity scores.

**Limitations:** Other cognitive mechanisms and white matter tracts may play a role in temporal impulsivity.

**Conclusions:** This study provides evidence that individual differences observed in impulsivity may be explained by variability in limbic fronto-striatal tracts. We emphasize the importance of investigating the spectrum of impulsivity in HD, less prevalent than other psychiatric features, but impacting the quality of life of patients and their caregivers.

## Introduction

Huntington’s disease (HD) is an autosomal-dominant, neurodegenerative disorder caused by an expansion of a CAG repeat in the *HTT* gene^1^. Degeneration results in atrophy of the basal ganglia and cerebral cortex and disruption of the cortico-striatal networks^2^. This leads to progressive motor and cognitive deficits and behavioral abnormalities^3^. Although depressive mood, apathy, perseverative behavior or irritability are the most extensively reported and studied neuropsychiatric symptoms in HD, problems related to poor impulse control and the consequent impulsivity are also recognizable in HD patients^4-6^. However, studies addressing impulsivity in HD are scarce and most of the available knowledge refers to its relationship with other neuropsychiatric features such as mania, irritability, and aggression^3,7^.

Impulsivity is a complex, multifaceted construct broadly defined as the tendency to act or react prematurely, without a plan or a estimation of the consequences of actions, easily leading to many failures of self-control including addictive and healthy risk behavior patterns^8^. From the different impulsivity domains, temporal impulsivity refers to the tendency to prefer smaller, immediate rewards to those that are larger, but delayed, even if the reward is worth the effort to wait^9^. There is a natural tendency to devaluate the reinforcement as the delay to receipt increases, opting for smaller, but immediate rewards over possible long-term higher-value outcomes. However, the reward system may also favor delayed rewards if the reward is worth the effort to wait by engaging cognitive control mechanisms that facilitate the inhibition of this urge.

There are widely ranging differences in the representation that people have of the valuation of rewards associated to delayed outcomes^10^. In particular, the delay has a stronger effect on subjective value for more impulsive behaviors and impulse control disorders^11–14^. That is, discount rates are steep even if the delayed discounts have high values. In addition, other motivational bias, such as enhanced sensitivity to reward (SR) and reduced sensitivity to punishment (SP), have been identified as potential modulating factors of the subjective reward valuation associated with impulsivity^15,16^.

Temporal impulsivity is typically studied through a Delay Discounting (DD) paradigm^17^. This task utilizes manipulation of smaller, immediate monetary rewards over larger, postponed rewards to assess decision-making abilities and specifically characterize the point at which the preference for an immediate reward changes to one that is delayed. High delay-discounting rates have been reported in populations with impulse control disorders, including pathological gamblers^11^, and patients with psychiatric disorders such as schizophrenia^12^, or neurodegenerative disorders such as Alzheimer’s^13^, among others.

To our knowledge, only two studies have used DD procedures to study temporal impulsivity in the context of HD both done in transgenic rat models of HD^18,19^. These studies demonstrated that transgenic HD rats presented higher levels of choice impulsivity and a lack of behavioral inhibition compared to control rats, translating to poor efficiency in gambling tasks and steeper DD curves. As such, the delay degrades the value of reward faster, resulting in the preference of smaller, more immediate rewards.

Different neuroimaging studies have investigated the neural systems involved in DD^20^. Aligning functional neuroimaging studies have demonstrated the involvement of the ventral striatum and the ventromedial prefrontal cortex in the explanation of the variability in DD^10,21^. However, little is known about the influence of the ventral cotico-striatral tracts that connect the aforementioned target regions, including the uncinate fasciculus tract (UF) and the accumbofrontal tract (NAcc-OFC), in individual differences of temporal impulsivity. More specifically, no study has explored this relationship in HD gene-mutation carriers, and the effect of the disease on ventral pathways such as the NAcc-OFC has not been investigated yet.

UF, a bidirectional limbic fiber pathway, connects the anterior temporal lobe with the orbitofrontal cortex (OFC) and the amygdala. the UF plays a role in the functional relationship between the prefrontal cortex and the mesial temporal lobe structures, and has been related with emotional and reward processing, i.e. reward-based decisions^22,23^.

The NAcc-OFC tract projects from the prefrontal cortex to the NAcc. Structural changes to the prefrontal cortex and striatum in humans has been associated with impulsive behavior^24^. In addition, a recent study^25^ also observed a significant positive association with impulsivity and white matter (WM) integrity of the NAcc-OFC.

Through this framework, the aim of the present study is to investigate the behavioral and neural bases underlying the individual differences in impulsivity in HD as evaluated by a DD task. To this end, we explore the relationship between levels of impulsivity and WM microstructure of the NAcc-OFC and the UF, two major ventral fronto-striatal fiber bundles associated with reward processing.

## Methods

### Participants

Forty-seven HD gene-expansion carriers and 36 healthy controls matched for age, sex and years of education participated in this study. Participant demographic and clinical information are detailed in Table 1. HD participants were all confirmed gene-mutation carriers with ≥36CAG repeats. Because neural degeneration, and the resulting cognitive and psychiatric symptoms, are often present long before the clinical diagnosis of HD^3^, we studied the disease as a continuum across manifest and pre-manifest individuals, unless otherwise specified.

**Table 1.** Sociodemographic and clinical characteristics of study participants.

|  | Control | Manifest | Pre-manifest |
| --- | --- | --- | --- |
| ***N*†** | 35 | 25 | 22 |
| **Sex (f/m)** | 18/17 | 14/11 | 18/4 |
| **Age (years)** | 44.00 ± 11 | 50.80 ± 9.39 | 36.64 ± 8.7 |
| **Education (years)** | 12.74 ± 2.7 | 11.24 ± 3.27 | 13.72 ± 2.81; N = 22 |
| **UHDRS-TMS** | — | 21.52 ± 12.58; N=25 | 1.52 ± 3.0; N = 21 |
| **UHDRS-cogscore** | — | 189.65 ± 59.48; N=23 | 297.31 ± 62.54; N = 22 |
| **CAP** | — | 113.02 ± 20.71 | 78.71 ± 14.57 |
| **DD (Impulsivity)** | -0.22 ± 0.56; N= 33 | 0.37 ± 1.35; N=17 | 0.06 ± 1.18; N=18 |
| **SR** | -0.11 ± 0.92; N= 34 | 0.21 ± 0.96; N= 24 | -0.07 ± 1.18; N=19 |
| **SP** | 0.06 ± 0.97; N= 33 | -0.058 ± 1.12; N= 24 | -0.02 ± 0.94; N19 |

Data presented as mean ± standard deviation. † Number of participants listed in individual cells when differing from this number for each group. CAP=Standardized CAG-Age Product; DD=Delay discounting, the Overall-k from Spanish version of the original Kirby Delay-Discounting Rate Monetary Choice Questionnaire; f=females; m=males; *N*=number of participants; SR=Sensitivity to Reward from the Spanish version of the Sensitivity to Punishment and Sensitivity to Reward Questionnaire; SP=Sensitivity to Punishment from the Spanish version of the Sensitivity to Punishment and Sensitivity to Reward Questionnaire; UHDRS-cogscore=Unified Huntington's Disease Rating Scale total cognitive score; UHDRS-TMS=Unified Huntington's Disease Rating Scale total motor score (Huntington Study Group, 1996).

Twenty-five of the gene-mutation carriers were manifest HD patients, defined as those with a diagnostic confidence level (DCL)≥4 on the Unified Huntington’s Disease Rating Scale (UHDRS). Eleven of the gene-mutation carriers were pre-manifest participants with a DCS<4. One control and two HD participants did not receive a diffusion-weighted image scan due to claustrophobia. Furthermore, WM microstructural outliers were identified and removed as Z-scores greater than |2.5|. The specific *N* is detailed for each test. None of the participants reported previous history of neurological disorder other than HD. The study was approved by the ethics committee of Bellvitge Hospital in accordance with the Helsinki Declaration of 1975. All participants signed a written declaration of informed consent.

### Clinical evaluation

The HD group underwent the clinical-UHDRS evaluation, which comprises motor, cognitive and behavioral subscales. The standardized CAG-Age Product (CAP) score was computed as CAP=100×age×(CAG–35.5)/627^26^. CAP has been used to model the effects of age and CAG length on various measures of the HD state and is assumed to reflect the effects of lifelong exposure to mutant huntingtin protein. Neurologists or neuropsychologists specialized in movement disorders carried out all clinical assessments.

### Questionnaire Measures

#### Delay-Discounting Task

In order to assess the individual rate of discounting reward value by delay, a Spanish version of the original Kirby Delay-Discounting Rate Monetary Choice Questionnaire was administrated on paper. For each of the 30 items, the participant had to choose between two options, a smaller reward that the participant could receive sooner (immediate) or a larger amount that could be received in the future (delayed), based on a monetary choice. Additionally, three control items that detailed higher amounts at present compared to smaller amounts in the future were included. The rate of discounting (*k-*score) is the slope of the hyperbolic function through the individual subjective value of delayed rewards and was automatically scored^27^

### Sensitivity to Punishment and Sensitivity to Reward Questionnaire

A Spanish version of the Sensitivity to Punishment and Reward Questionnaire (SPSRQ) was completed by participants. The SPSRQ is a self-report questionnaire with 48 Yes/No response items composed of two scales: SP and SR. The score of each subscale ranges from 0–24, with higher scores indicating a higher SP (selective responsiveness to fear and anxiety-evoking stimuli) or SR (selective responsiveness to stimuli with emotional well-being, reward, and other consummatory behaviors).

### MRI data acquisition

MRI data were acquired using a 3T whole-body MRI scanner (Siemens Magnetom Trio; Hospital Clínic, Barcelona), through a 32-channel phased array head coil. Structural images comprised a conventional high-resolution 3D T1-image (magnetization-prepared rapid-acquisition gradient echo sequence (MPRANGE), 208 sagittal slices, repetition time (TR)=1970ms, echo time (TE)=2.34ms, inversion time (TI)=1050ms, flip angle=9º, field of view (FOV)=256mm, 1mm isotropic voxel).

Diffusion weighted MRI data were acquired using a dual spin-echo diffusion-tensor imaging (DTI) sequence with GRAPPA (reduction factor of 4) cardiac gating, with TE=92ms, 2mm isotropic voxels, no gap, 60 axial slices, FOV=23.6cm. In order to obtain the diffusion tensors, diffusion was measured along 64 noncollinear directions, using a single b-value of 1500s/mm^2^ interleaved with 9 non-diffusion (b=0) images. In order to avoid chemical shift artifacts, frequency-selective fat saturation was used to suppress fat signal.

### DW-MRI tractography analysis

The gold-standard automated probabilistic tractography approach, Tracts Constrained by UnderLying Anatomy (TRACULA)^28^, was utilized for the dissection of the UF. Because the NAcc-OFC is not included in the TRACULA atlas, we employed a deterministic dissection approach using TrackVis for the NAcc-OFC.

### Preprocessing of DTI data

For the UF, DTI were automatically processed using FreeSurfer v6.0 software (http://surfer.nmr.mgh.harvard.edu/). Specifically, head motion and eddy-current correction were first performed using the FMRIB’s Diffusion Toolbox in FMRIB’s Software Library (FSL, http://www.fmrib.ox.ac.uk/fsl/fdt) and the gradient matrix was rotated accordingly^29^. Boundary-based registration method was used for the affine intra-subject alignment between the diffusion-weighted and anatomical images, and to an MNI152 template^30^. The diffusion tensor was then reconstructed using a standard least squares tensor estimation algorithm for each voxel and then fractional anisotropy (FA), mean diffusivity (MD), and radial diffusivity (RD) maps were calculated.

With regard to the NAcc-OFC, fiber orientation distributions for the NAcc-OFC were reconstructed using a spherical deconvolution approach based on the damped version of the Richardson-Lucy algorithm implemented in StarTrack software (http://www.natbrainlab.co.uk). In particular, a combination of spherical deconvolution parameters was selected to resolve crossing and avoid spurious peaks in gray matter or cerebral spinal fluid (fixed fiber response corresponding to a shape factor of α=2×10– 3mm2/s; 200 algorithm iterations, regularization threshold [=0.04 and regularization geometric parameter v=8) (Dell’Acqua et al. (2010), for further details).

Whole-brain tractography of the NAcc-OFC was next performed using a b-spline interpolation of the diffusion tensor field and Euler integration to propagate streamlines following the directions of the principal eigenvector with a step size of 0.5mm. Tractography was stopped when FA<0.2 or when the angle between two consecutive tractography steps was larger than 35°. Finally, tractography data and diffusion tensor maps were exported into TrackVis (http://www.trackvis.org) for manual dissection of the tracts.

### Tractography dissections

Virtual *in vivo* DTI reconstruction of the UF tract were carried out bilaterally using TRACULA^31^. TRACULA is the gold-standard approach for the probability reconstruction of the main WM brain pathways and has been previously validated in HD patients. Briefly, to estimate the posterior probability distribution, the algorithm combines the individual participant’s local diffusion orientations extracted from the ball-and-stick model of diffusion ^32^, with prior information of the pathways from a set of training participants where the tracts of interest were labeled manually.

Virtual *in vivo* DTI dissections of the NAcc-OFC were carried out bilaterally in native space FA maps using two-regions of interest (ROI) approach. To dissect the NAcc-OFC, fibers projecting freely from the NAcc ROI were selected and restricted to terminate in the OFC, defined following previously proposed anatomical guidelines^33^. The NAcc ROIs were segmented by using the FSL FIRST toolbox and then registered to the individual native diffusion space using the FSL FLIRT^34^ and FNIRT^35^ modules after normalizing both the structural T1-images and FA maps.

Controls and patients were randomized for the tractography that was performed by one dissector to blind participant identity. FA, MD and RD values were extracted for analysis of WM microstructure.

### Statistical analyses

Statistical analyses were performed in SPSS (v.25, SPSS Inc, Chicago, USA). Independent two-tailed t-tests and Cohen’s d were applied. Both were used to describe differences between HD and control groups in temporal impulsivity (*k*-score), motivational SR and SP as well as WM disturbance within dissected tracts. Then, in order to further analyze differences between temporal impulsivity between groups, both HD patients and controls were categorized into quartiles according to their raw *k*-score. We compared the top quartile (i.e. highest impulsivity) between both groups. For tract integrity, we also compared differences within the HD group using the top and lower quartiles. In order to investigate the relationship between impulsivity and HD progression, we employed a one-way ANOVA with post-hoc comparisons, taking into account controls, pre-manifest and manifest groups. In addition, Pearson’s correlations were used to assess a possible association between impulsivity scores and CAP, a proxy of disease stage.

We investigated differences in structural connectivity between HD and controls in the selected tracts by using independent two-tailed t-tests. Specifically, WM disturbances were estimated from extracted FA, MD and RD mean values of the UF and NAcc-OFC tract bilaterally.

Afterwards, we explored whether differences in WM microstructure in the main tracts of interest may explain variability in impulsivity in HD participants. An initial multiple linear regression analysis was performed, including temporal impulsivity as a dependent variable with the predictor variables being one of the diffusion indices (FA, MD, or RD) of the two tracts of interest. Sex and CAP were included as control variables. The same process was repeated for each diffusion index. A second multiple regression analysis was used to further explore potential motivation bias in temporal impulsivity by additionally including SR and SP as predictor variables. Finally, the same regression models were performed on the control group and again using all the participants as a whole (HD and control group combined). This was done in order to investigate whether the observed effects were specific to HD participants, or instead explained the variability of impulsivity and WM microstructure in the general population. The false discovery rate (FDR) was used to correct all t-tests and correlations for multiple comparisons (q=0.05). We controlled for four comparisons (two tracts × two hemispheres) when structural connectivity effects between groups were investigated. We also controlled for two comparisons (two models) in the regression analysis. Both raw p-values (P) and the p-adjusted FDR values (P-adj) are reported.

## Results

### Behavioral results

When comparing the impulsivity scores between HD patients (pre-manifest and manifest combined) and controls, we found HD patients displayed a trend toward higher impulsivity scores (*t*(47.44)=-1.83, *P*=.074, two-tail, Cohen’s *d*=.094). In order to further study the distribution of impulsivity, the impulsivity scores of the top quartile were compared against groups. We observed that the highest impulsivity patients had higher statistically significant scores compared to the highest impulsivity controls (*t*(12.11)=2.23, *P*=.045, two-tail, Cohen’s *d*=1.01). The DD task showed excellent internal consistency between participants. Specifically, the overall task consistency of the control group was of 96,84%, while the overall consistency of the HD was of 94,30%, providing convergent evidence that the participants were not engaged in random responding nor they had unexpected response patterns.

In addition, we examined potential differences in impulsivity scores between the three groups (pre-manifest, manifest and control) using a one-way ANOVA with post-hoc analyses. No significant between-group differences were found [(F(2.65)=2.05, *P*=.137)]. Likewise, no significant correlations were found when examining the association between impulsivity and CAP (*r*=-.03, *P*=0.87). No significant differences were found between SR (*t*(71.07)=.42, *P*=.68, *d*=.097) or SP (*t*(74.32)=-.89, *P*=.38, *d*=.78) when comparing the HD and the control group. For details, see Table 1 and Figure 1A.

**Figure 1.**
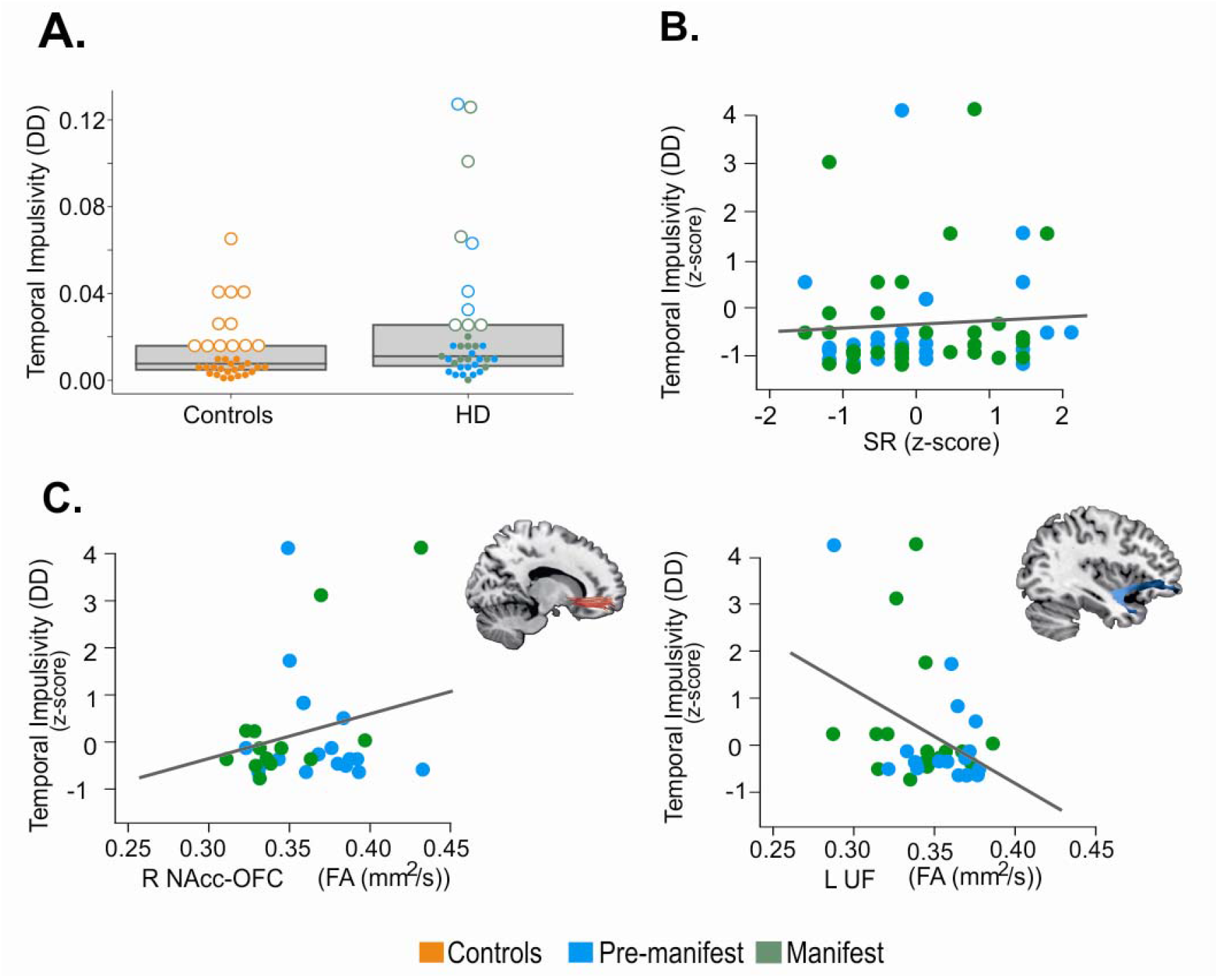
Behavioral and microstructural modulations of temporal impulsivity in Huntington’s disease. (**A**) Impulsivity scores between the controls and HD group: Scatterplot of behavioral correlates displaying the impulsivity scores between HD patients (pre-manifest and manifest combined) and controls. Box plots indicate the first, second (median), and third quartile limits. Individuals in the fourth quartile are marked with an empty circle, and individuals in other quartiles are marked with solid dots. (**B**) Displays the relationship between impulsivity levels and SR. (**C**) Displays the relationship between impulsivity levels and white matter disturbance in the two tracts of interest. The two scatterplots display the significant association obtained in the multiple regression model between impulsivity and FA in the right accumbofrontal fasciculus (NAcc-OFC), and left uncinate fasciculus (UF). Lower FA values indicate more severe damage to white matter microstructure. Linear regression line is fit for illustration in each scatterplot. DD = delay-discounting, temporal impulsivity scores; FA = fractional anisotropy; NAcc-OFC = Accumbofrontal tract; SR = Sensitivity to reward; UF = uncinate fasciculus.

### Differences in structural connectivity between HD and controls

The HD group revealed higher mean RD in both the UF and NAcc-OFC tracts when compared with controls, demonstrating abnormal WM microstructure (Table 2). Specifically, there was a significant increase in RD values in HD individuals compared with controls in the left NAcc-OFC (*t*(70.83)=-2.20, *P*=.031, *P-adj*=.034, *d*=.51) and left UF (*t*(73.87)=-2.24, *p*=.03, *P-adj*=.03, *d*=.51) as well as the right UF (*t*(75.74)=-2.69, *P*=.01, *P-adj*=0.03, *d*=.59). Moreover, HD patients showed reduced connectivity in the right UF when compared with controls. In particular, there was a significant increase in MD values in the right UF (*t*(75.99)=-2.16, *p*=.03, *P-adj*=.03, *d*=.48) in the HD group compared with controls (four comparisons), in addition to a significant decrease in the FA values of the right UF (*t*(75.20)=2.60, *P*=.01, *P-adj*=.03, *d*=.58) (four comparisons).

**Table 2.** Mean FA and MD diffusion values in manifest Huntington’s disease patients and controls.

|  |  |  | HD | *N* | Controls | *N* | *t* | *P*-value |
| --- | --- | --- | --- | --- | --- | --- | --- | --- |
| NAcc-OFC | Left | FA | -0.057 ± 1.082 | 40 | 0.069 ± 0.903 | 33 | 0.539 | 0.592 |
|  |  | MD | 0.194 ± 1.120 | 40 | -0.235 ± 0.786 | 33 | -1.918 | 0.059 |
|  |  | RD | 0.223 ± 1.074 | 40 | -0.270 ± 0.841 | 33 | -2.201 | 0.031* |
|  | Right | FA | -0.039 ± 1.016 | 37 | 0.051 ± 0.995 | 28 | 0.361 | 0.719 |
|  |  | MD | 0.189 ± 1.045 | 37 | -0.259 ± 0.889 | 27 | -1.850 | 0.069 |
|  |  | RD | 0.192 ± 1.006 | 37 | -0.254 ± 0.950 | 28 | -1.831 | 0.072 |
| UF | Left | FA | -0.190 ± 0.904 | 45 | 0.260 ± 1.078 | 33 | 1.947 | 0.056 |
|  |  | MD | 0.164 ± 1.095 | 45 | -0.224 ± 0.818 | 33 | -1.789 | 0.077 |
|  |  | RD | 0.210 ± 1.042 | 44 | -0.280 ± 0.880 | 33 | -2.236 | 0.028* |
|  | Right | FA | -0.236 ± 1.046 | 45 | 0.322 ± 0.847 | 33 | 2.598 | 0.011* |
|  |  | MD | 0.196 ± 1.089 | 45 | -0.267 ± 0.805 | 33 | -2.161 | 0.034* |
|  |  | RD | 0.238 ± 1.095 | 45 | -0.325 ± 0.753 | 33 | -2.688 | 0.009* |

Subsequently, when comparing impulsivity scores in the top quartile (i.e. highest impulsivity), significant differences were seen between HD individuals and controls (*t*(12.11)=-2.23, *P*=.045). The analysis of quartiles was also used to study tract integrity within HD group. In this case, when comparing the top with the lowest impulsivity quartiles of HD individuals, the FA of the left UF (*t*(13.88)=2.96, *P*=.01, *P-adj*=.02) and left NAcc-OFC (*t*(6.31)=2.52, *P*=.043, *P-adj*=.043) were the only with significant differences between the two extreme quartiles.

### Multiple linear regression models

Multiple linear regression analyses were performed to investigate the association between temporal impulsivity and the tracts of interest in HD participants. Specifically, the first regression model revealed that increased WM integrity (higher FA), of the right NAcc-OFC (β=0.64, *P*=.016) and decreased WM integrity (lower FA) of the left UF (β=-1.10, *P=*.03) were significant predictor variables of more impulsive behavior, explaining approximately half of the variance of the impulsivity score (R^2^=.44, F(6.22)=2.92, *P*=.03, *P*-*adj*=.03). Interestingly, the second model, which additionally included SR and SP scores as predictors, was found to more completely explain the variance of temporal impulsivity scores (R2= .64, F(8.18)=4.16, P=.01, P-adj=.02) (Figure 1B). This revealed SR (β=.65, P=.02) to be a significant contributor to the model (Figure 1C), in addition to the right NAcc-OFC FA (β=.96, P=.001) and left UF FA (β=-1.59, P=.01). Furthermore, when the multiple regression models were repeated in the control group and again in the HD and control combined groups, no regression coefficients were found to be significant.

## Discussion

We aimed to investigate the spectrum of DD behavior as a proxy of temporal impulsivity in individuals with HD. We observed individual differences in the distribution of temporal impulsivity among patients. Particularly, a subset of pre-manifest and manifest HD participants exhibited higher temporal impulsivity scores than controls. The fact that these effects were observed only in select individuals is in accordance with previous studies, in which a lower overall prevalence of impulsivity is observed in comparison to other neuropsychiatric symptoms^3,6,36^. Nevertheless, the true prevalence of impulsivity may not be fully characterized, as this symptom is not usually evaluated as temporal impulsivity, neither as an isolated entity in neuropsychiatric assessments, but rather encompassed into other neuropsychiatric symptoms or characterized as disinhibition, irritability, or addictive behaviors.

The HD group exhibited disturbed microstructure in the main tracts of interest (UF bilaterally, and the left NAcc-OFC) when compared to controls, represented by higher RD, MD, and lower FA. Histologically, an increase in RD represents a biomarker of myelin abnormalities, while increased MD accompanied with reduced FA might represent a sign of loss tissue architecture. WM abnormalities in the UF have already been shown in both premanifest and manifest HD patients^37^. However, to our knowledge, this is the first study to show altered structural connectivity in the NAcc-OFC tract in HD. This is a very interesting result since the gradients of atrophy in HD start at dorsal regions of the caudate and putamen and progress to lateral regions, leaving the ventral striatum relatively spared until late stages of the disease.

Neurobiologically, individual differences in levels of temporal impulsivity were explained by variability in WM microstructure. Our results revealed increased structural connectivity (higher FA) in the NAcc-OFC, predicting high values of temporal impulsivity. This aligns with previous HD studies, where WM integrity of the NAcc-OFC showed a significant positive association with temporal impulsivity in healthy populations^25^. Likewise, the UF has also been implicated with emotion-based evaluative processes, cognitive flexibility and impulse control in different disorders^23,38^. Moreover, in the present study, reduced structural connectivity (lower FA) was found to be associated with higher temporal impulsivity scores. In the same line, this conflicts with previous studies in delay gratification, showed that where individual variability in the UF microstructure was found to predict the ability to control the temptation of an immediate reward in order to receive a larger, but delayed reward. For example, in healthy individuals, greater tolerance for delayed rewards, as measured by DD, predicted increased FA of the UF region in both hemispheres^39^. Simultaneously, these inverse connectivity patterns (NAcc-OFC with increased FAvs.UF with decreased FA) align with prior studies, which have consistently shown that parallel limbic ventral cortico-striatal networks may operate in a reciprocal manner^24,40^. The NAcc-OFC and UF circuits may accordingly engage two different mechanisms, where reward and cognitive control processes guide temporal impulsivity^10,21,40^. Thus, the reward system functions to obtain immediate rewards by estimating the incentive value, while the cognitive control system inhibits this response by predicting the consequences of the delay and/or the possible outcome^10^. This supports the hypothesis that these two processes operate in tandem, each represented by a parallel limbic ventral cortico-striatal network^24,40,41^.

Additionally including SR as a predictor factor revealed a mediator role between SR and temporal impulsivity, exemplifying that individuals with heightened SR may be driven to elicit greater impulsivity^8,40^. Specifically, SR is regulated by the behavioral activation system, which influences the extent to which behavior is motivated by reward-relevant stimuli. Hence, heightened SR is linked with addiction and risk-taking behaviors in both general and clinical populations^42^. In HD individuals, the present results likewise reveal that the reward valuation decreased with greater delay. This translates to a tendency to prefer immediate over delayed monetary rewards, highlighting the value of the reward above the possible future outcomes. Such enhanced allure of the reward aspect even in risky or uncertain outcomes can manifest as impulsive decision-making or a higher predisposition to develop a gambling addiction, as has been suggested in HD^42,43^.

This study shares certain limitations. It is important to highlight that the study was exploratory in nature, aiming to break a new ground in the study of temporal impulsivity in HD through the use of the DD task, as potentially modulated by an interplay between reward and cognitive control networks. Additionally, other cortico-striatal frontal tracts beyond the NAcc-OFC and UF may play a role in temporal impulsivity. Similarly, it is possible that other factors, such as time perception, might mediate impulsive behavior in DD tasks^44–46^. Moreover, examining the relationship between impulsivity and apathy, which has been cited as being both positively and negatively related in other contexts^47^, could clarify the multidimensional nature of underlying neuropsychiatric constructs. The continuation of this research line in a larger sample size and distinct patient populations may serve to elucidate these questions.

Overall, these findings highlight the importance of further exploring the heterogeneous psychiatric symptomatology in HD. This includes those disease features that may be less prevalent, but still bear a significant burden on patient and caregiver quality of life, thus opening a door to more personalized care.

## Acknowledgments

We are grateful to the patients and their families for their participation in this project. We would also like to thank Dr. Saül Martinez-Horta, Dr. Jesús Pérez Pérez, Dr. Jaime Kulisevksy; Pilar Sanchez, Dr. Esteban Muñoz (Hospital Clínic) for help with clinical evaluation.

## Financial Disclosures of all authors for the previous 12 months

The authors declare that there are no additional disclosures to report.

## Data availability

The raw data that support the findings of this study are available from the corresponding author upon reasonable request after approval of local institutional review board.

## Ethical Compliance Statement

The study was approved by the ethics committee of Bellvitge Hospital in accordance with the Helsinki Declaration of 1975.

All participants signed a written declaration of informed consent.

We confirm that we have read the Journal’s position on issues involved in ethical publication and affirm that this work is consistent with those guidelines.

